# Tracking temporal hazard in the human electroencephalogram using a forward encoding model

**DOI:** 10.1101/233551

**Authors:** Sophie K. Herbst, Lorenz Fiedler, Jonas Obleser

**Affiliations:** Department of Psychology, University of Lübeck Ratzeburger Allee 160, 23552 Lübeck, Germany

## Abstract

Human observers automatically extract temporal contingencies from the environment and predict the onset of future events. Temporal predictions are modelled by the hazard function, which describes the instantaneous probability for an event to occur given it has not occurred yet. Here, we tackle the question of whether and how the human brain tracks continuous temporal hazard on a moment-to-moment basis, and how flexibly it adjusts to strictly implicit variations in the hazard function. We applied an encoding-model approach to human electroencephalographic (EEG) data recorded during a pitch-discrimination task, in which we implicitly manipulated temporal predictability of the target tones by varying the interval between cue and target tone (the foreperiod). Critically, temporal predictability was either solely driven by the passage of time (resulting in a monotonic hazard function), or was modulated to increase at intermediate foreperiods (resulting in a modulated hazard function with a peak at the intermediate foreperiod). Forward encoding models trained to predict the recorded EEG signal from different temporal hazard functions were able to distinguish between experimental conditions, showing that implicit variations of temporal hazard bear tractable signatures in the human electroencephalogram. Notably, this tracking signal was reconstructed best from the supplementary motor area (SMA), underlining this area’s link to cognitive processing of time. Our results underline the relevance of temporal hazard to cognitive processing, and show that the predictive accuracy of the encoding-model approach can be utilised to track abstract time-resolved stimuli.

**Significance Statement:** Extracting temporal predictions from sensory input allows to process future input more efficiently and to prepare responses in time. In mathematical terms, temporal predictions can be described by the hazard function, modelling the probability of an event to occur over time. Here, we show that the human EEG tracks temporal hazard in an implicit foreperiod paradigm. Forward encoding models trained to predict the recorded EEG signal from different temporal-hazard functions were able to distinguish between experimental conditions that differed in their build-up of hazard over time. These neural signatures of tracking temporal hazard converge with the extant literature on temporal processing and provide new evidence that the supplementary motor area tracks hazard under strictly implicit timing conditions.

## Introduction

Time provides the structure of our experience, and is the basis of many cognitive processes, like motor acts and speech processing. Even when we do not consciously track the passage of time, the extraction of temporal contingencies from the environment allows us to generate temporal predictions about the occurrence of future events (for reviews see Coull, 2009; Nobre et al., 2007).

In mathematical terms, temporal predictions can be expressed by the hazard function, which models for any time point in a pre-defined interval the conditional probability of an event to occur, given it has not yet occurred. Crucially, temporal hazard depends on both, elapsed time in the current situation and the observer’s expectation about possible durations, that is the underlying distribution of durations. For example, when waiting at a traffic light, one might assume that waiting times vary in the range of several seconds, and that any duration in that range is equally likely (i.e. assuming a uniform distribution of durations). In this case the hazard for the light to turn green rises monotonically with elapsed time. Or, one could assume that some durations are more probable than others, for example because most traffic lights change from red to green at around 30 s (i.e. assume a distribution with one or more peaks resulting in a modulated hazard function). Little is known yet about how flexibly we extract statistical distributions of durations and utilize the resulting hazard functions to create temporal predictions, and which cognitive and neural processes are involved in this process.

Previous studies have shown that temporal hazard shapes behavioral responses (Karlin, 1966; Niemi and Näätänen, 1981; Tomassini et al., 2016; Tsunoda and Kakei, 2011) and is reflected in neural processing in monkeys (Akkal et al., 2004; Janssen and Shadlen, 2005; Jazayeri and Shadlen, 2015) and humans (Bueti et al., 2010; Cravo et al., 2011; Cui et al., 2009; Jazayeri and Shadlen, 2015; Trillenberg et al., 2000; Praamstra et al., 2006). For instance, Janssen und Shadlen (2005) tested whether monkeys learn to distinguish a unimodal from a bimodal duration distribution and showed that reaction times and recordings of single-neuron activity in the lateral intra-parietal area correlated with the respective hazard functions of unimodal or bimodal duration distributions. In humans, Bueti et al. (2010, using fMRI) and Trillenberg et al., (2000, using EEG), showed that neural activity prior to target onset correlated with the respective hazard function. These studies used a so-called ‘set-go-task’, in which the participant has to withhold a response from a set-cue until the presentation of a go-cue, a task which enforces the use of temporal hazard. Trillenberg et al.’s participants were even informed about the underlying foreperiod probability distributions, which might have promoted the use of explicit timing strategies.

Here, our aim was to test whether observers utilize temporal hazard in a sensory task with no explicit incentives for timing. In contrast to explicit timing situations, in which participants are asked to provide an overt estimate of elapsed time, implicit timing does not require an overt judgement of time, but assumes that temporal contingencies guide response behaviour in a covered way (Coull and Nobre, 2008; Nobre and van Ede, 2018). Building on the earlier work, we were interested in whether human observers can distinguish between different levels of implicit temporal predictability. Instead of using duration distributions that differ with respect to the most likely time point at which an event is predicted to happen, we used three different unimodal probability distributions, all with equal mean. To vary the level of temporal predictability, we used either a uniform distribution (non-predictive; mean duration = 1.8 s), or two Gaussian shaped distributions with the same mean, and a larger (weakly predictive) and smaller (strongly predictive) standard deviation (see Figure 1). Previously, we showed that the non-predictive and strongly predictive conditions evoked distinguishable correlates of temporally predictive processing (Herbst and Obleser, 2017). Here, we asked whether the concept of temporal hazard, derived from the foreperiod distributions can explain human behaviour in an implicit foreperiod paradigm, and whether it bears tractable signatures in the human EEG.

**Figure 1:**
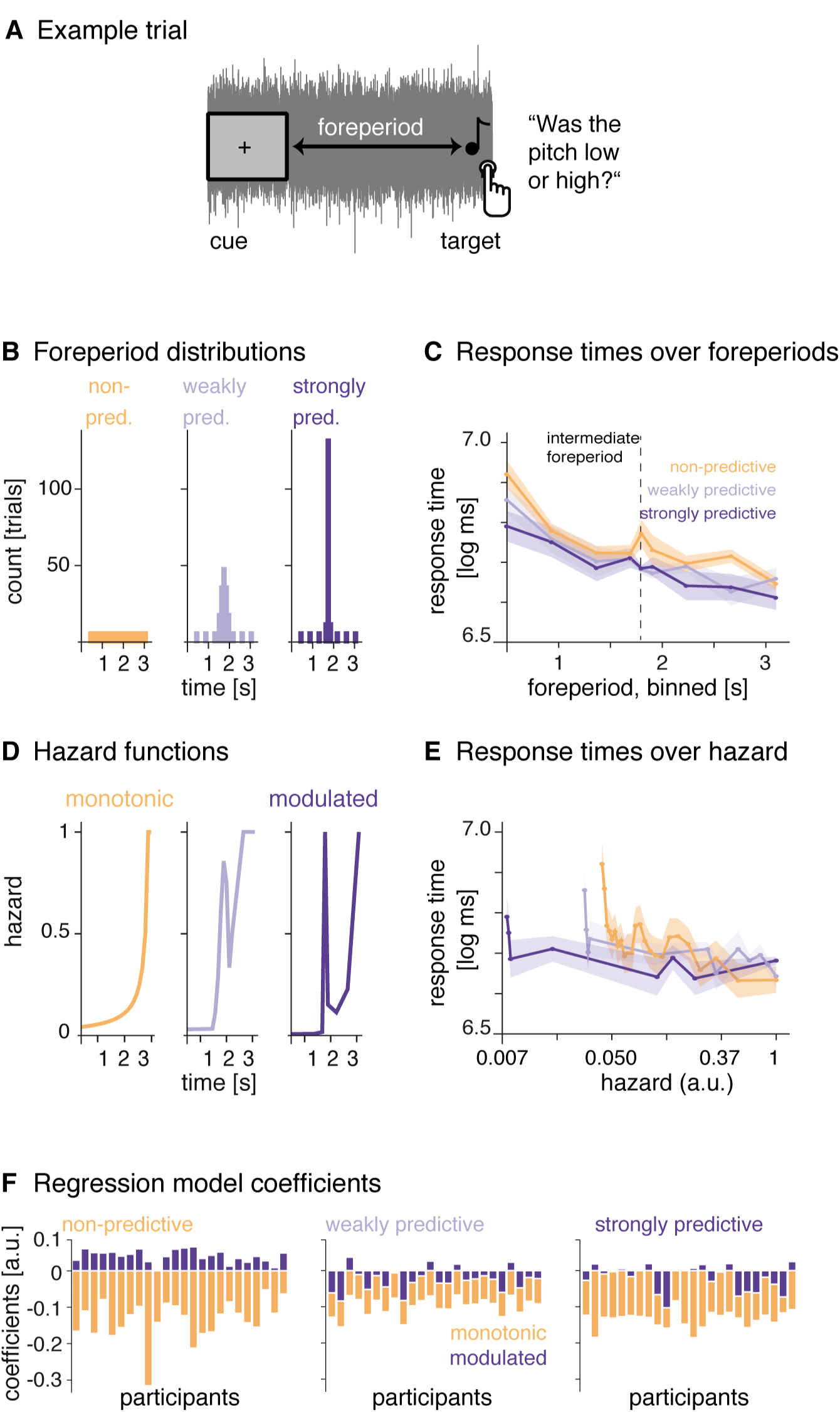
Paradigm and response times. A: Example trial. On each trial, the simultaneous onset of the fixation cross and the noise served as a temporal cue. After a variable foreperiod interval, a single target tone was then presented and participants had to judge its pitch as low or high. B: Foreperiod probability distributions. Foreperiods for each block were drawn from one out of three distributions: a uniform distribution (non-predictive), and two normal distributions with larger and smaller standard deviations (weakly and strongly predictive). C: Response times over foreperiods (binned): response times were longer at short foreperiods and shorter at long foreperiods. D: Hazard Functions resulting from the foreperiod probability distributions. E: Response times over temporal hazard (i.e. the value of the hazard function at the time of target onset, on a log-spaced axis): response times were longest at the lowest hazard. Shaded surfaces reflect standard error of the mean. F: relative regression coefficients per participant obtained by modelling response times per condition (panels from left to right) with the monotonic and modulated hazard functions as regressors.

For the uniform foreperiod distribution, the hazard function rises monotonically towards the end of the range of possible intervals, but for the two Gaussian distributions it is modulated and contains an earlier peak, too. To track the neural processing of temporal hazard over time, we measured response times and recorded neural activity with the electroencephalogram (EEG). We applied a forward encoding model approach (following Lalor et al., 2006, 2009; see also Fiedler et al., 2017; O’sullivan et al., 2015), using the hazard functions as time-resolved regressors to model the time domain EEG data. To our knowledge, it is the first time that the encoding model approach is applied to track the processing of an entirely abstract stimulus like temporal hazard, which is not related to the acoustic signal. Hence, a secondary aim of the presented approach was to proof the applicability of encoding models to track the processing of abstract stimuli in human time domain EEG. Assessing the fit between modelled and measured time domain EEG signals allowed us to quantify the representation of temporal hazard in the EEG, showing that the human brain distinguishes between different conditions of implicit temporal hazard.

## Methods

### Participants

A total of 24 healthy participants were tested (mean age 23.9 ± SD 2.2 years; 11 female, all right-handed), all reporting normal hearing and no history of neurological disorders. Participants gave informed consent and received payment for the experimental time (7 e per hour). The study procedure was approved by the local ethics committee (University of Leipzig, Germany). Note that this manuscripts reports extensive modelling and re-analysis of data that had been acquired for a study published earlier (Experiment II in Herbst and Obleser, 2017).

### Code and data accessibility

Publicly available Matlab toolboxes were used for the analyses (Fieldtrip, mTRF Toolbox: Crosse et al., 2016). The data and costum written analyses scripts are available at the Open Science Framework: https://osf.io/qbhma/

### Stimuli and paradigm

The EEG experiment was conducted in an electrically shielded sound-attenuated EEG booth. Stimulus presentation and collection of behavioral responses was achieved using the Psychophysics Toolbox (Brainard, 1997; Pelli, 1997). Responses were entered on a custom-built response box, using the fingers of the right hand for the pitch judgement and the fingers of the left hand for a subsequent confidence rating. Auditory stimuli were delivered via headphones (Sennheiser HD 25-SP II) at 50 dB above the individual’s sensation level. Stimuli were pure tones of varying frequencies (duration 50 ms with a 10 ms on- and offset ramp), embedded in lowpass (5 kHz) filtered white noise. Sensation level was individually predetermined for the noise using the method of limits with and tone-to-noise ratio was fixed at −16 dB. Target tones varied in individually predetermined steps around a 750-Hz standard, which was itself never presented. Participants had to perform a pitch discrimination task on a single tone presented in noise: ‘was the tone rather high or low?’ (for an exemplary trial see Figure 1A), followed by a confidence rating on a three level scale. The beginning of each trial was indicated by the simultaneous onset of the fixation cross and the noise. In 10% of all trials, participants received an additional, explicit timing question (‘Was the interval between the cue and the tone rather short or long?’) after the confidence rating. On average, 65% of these trials were answered correctly, showing that participants had processed the foreperiod duration and were able to retrieve it.

#### Foreperiod distributions and hazard functions

Foreperiods ranged from 0.5 to 3.1s (mean 1.8s) and were drawn from three different probability distributions (shown in Figure 1B). For the non-predictive condition, 25 discrete foreperiods were drawn from a uniform distribution. We call this condition non-predictive, because throughout the trial all tone onset times were equally likely. Nevertheless, due to the unidirectional nature of time, participants’ expectation for a target to occur should rise with elapsed time during these trials (as indicated by the hazard function, see below). For the two temporally predictive conditions, we drew foreperiods from two normal distributions with the same mean as the uniform distribution but varying standard deviations resulting in a weakly predictive condition (sd = 0.15; resulting in five discrete foreperiods) and a strongly predictive condition (sd = 0.05; resulting in three discrete foreperiods).

Furthermore, we added additional trials at three short (0.50, 0.93, 1.37 s) and and three long (2.23, 2.67, 3.10 s) foreperiods to both predictive conditions (see foreperiod distributions in Figure 1B), resulting in 25, 32 and 34 trials per block for the non-predictive, weakly predictive and strongly predictive conditions, respectively. The additional trials were included in all analyses, and foreperiods were presented in a counter-balanced manner. Inter-stimulus intervals (ISI) were drawn from a truncated exponential distribution (mean 1.5 s, truncated at 5s). This way we obtained maximally unpredictable ISI (Näätänen, 1971), in order to prevent entrainment to the stimulation over trials. To obtain the hazard functions (depicted in 1C), we transformed the discrete values of the foreperiod probability distributions (including the additional trials). The hazard function is defined as:

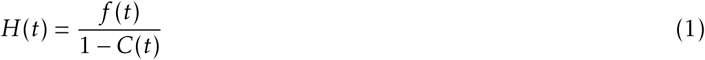

with t being time points throughout a trial t = 1 … T, f being the foreperiod probability distribution and C the cumulative function thereof. For normalization purposes, we first replaced infinite values that occurred for H(t) when C(t) = 1 by the maximum of the remaining values of H(f), and divided all values by that maximum to achieve hazard values ranging between 0 and 1. The uniform foreperiod distribution from the non-predictive condition results in a *monotonic hazard function* rising throughout the trial, indicating that if no tone has occurred yet, participants’ expectations for it to occur rises over time (Figure 1D, left). The weakly and strongly predictive conditions show the same rise towards the end, but additionally contain a peak at the intermediate foreperiod, due to the predictability manipulation (Figure 1D, middle, right). Out of the two predictable hazard functions, for the following analyses, we used only the hazard function from the strongly predictive condition. To obtain a maximal separation between the hazard functions from the non-predictive and strongly predictive conditions for the encoding models, we subtracted the monotonic from the modulated hazard function, thus removing the rise towards the end of the trial from the modulated hazard function and keeping only the peak, as the most distinctive feature. We refer to this hazard function as the *modulated hazard function*.

To be used as regressors in the encoding models fitted to the EEG data, the hazard functions had to match the sampling rate of the EEG data (200 Hz). To achieve this, we linearly interpolated the missing values. The resulting hazard functions are depicted Figure 2A.

**Figure 2:**
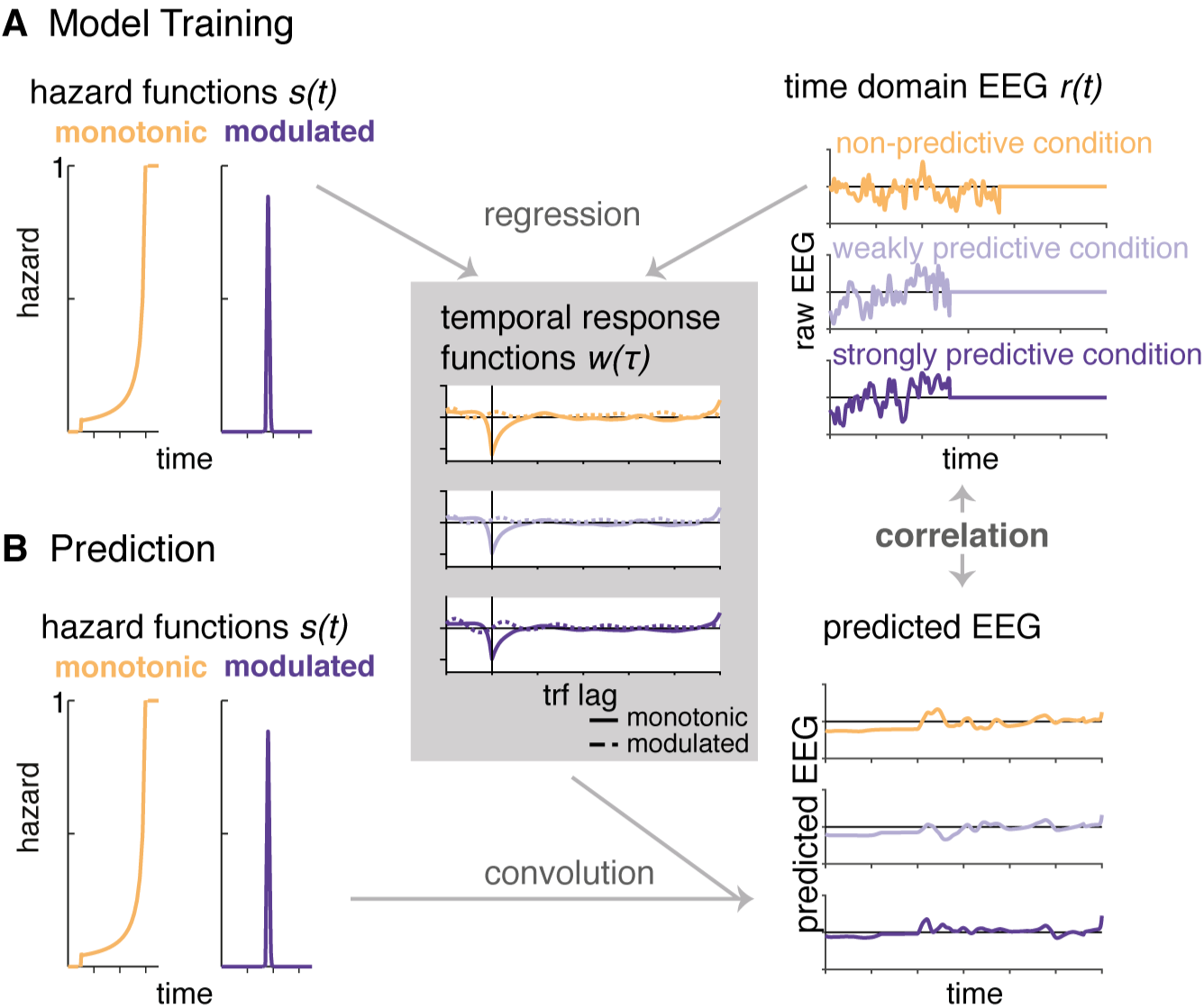
Schematic depiction of the encoding model approach. A: One encoding model was computed per condition, by regressing the two hazard functions (s in equation 2) on the time domain EEG data (r in equation 2; example trials are displayed, set to 0 at target onset. We obtained two temporal response functions (w in equation 2) per condition (right panel), one for the monotonic and one for the modulated hazard function. B: In a second step, we predicted EEG signals (example trials displayed in the bottom right panel) from each of the three models by convolving the hazard functions with the temporal response functions. Correlating the predicted and original EEG signals allowed us to test which model provides the best fit with the original EEG data.

## Procedure

At the beginning of the session, participants were briefed about the pitch discrimination task and the EEG recording. There was no mention of any aspects of timing prior to testing. After EEG preparation and the measurement of the individual sensation level, participants performed a training of 38 trials (25 for the pitch discrimination only plus 13 with the confidence rating added). During training, foreperiods were drawn from a uniform distribution to not induce any temporal predictions. Then, six condition blocks were presented in random order: each condition was presented once during the first and once during the second half of the experiment (three blocks per condition, one after the other), with random order between condition blocks, separated by breaks of self-determined length (60 s at minimum). There were 546 trials in total. After the experiment, participants were debriefed and asked whether they had detected the foreperiod manipulation. No participant spontaneously reported the predictability manipulation we applied. In a second question, the experimenter asked participants whether they detected any differences in the distributions of short and long foreperiods between blocks. Two participants stated they did detect a difference, which could be a hint that they noticed the predictability manipulation. In total, the experimental session lasted about 2.5 hours, including EEG preparation.

### EEG recording and pre-processing

Electroencephalographic data were continuously acquired from 68 electrodes (Ag–AgCl), including 61 scalp electrodes (Waveguard, ANT Neuro), one nose electrode, and two mastoid electrodes. The electrooculogram was acquired to record eye movements, with two electrodes placed horizontally to each eye and one placed vertically to the right eye. A ground electrode was placed at the sternum. All impedances were set below 10 kΩ. The nose electrode served as reference during recording. The data were acquired with a sampling rate of 500 Hz and a hardware-implemented passband of DC to 135 Hz (TMS International). EEG data were analysed using the Fieldtrip software package (versions 20160620, 20170501) for Matlab (MATLAB 2016a, 2017a). First, we applied high- and low-pass filtering to the continuous data using filters from the firfilt plugin (Widmann et al., 2015). The high-pass filter was a causal (one-pass-minphase) firws filter at 0.1 Hz with a transition bandwidth of 0.2 Hz. The lowpass-filter was a causal firws (one-pass, zero-phase) filter, with a 100 Hz cut-off and 3 Hz transition bandwidth. After filtering, data were re-referenced to linked mastoids, down-sampled to 200 Hz and demeaned. Next, the data were epoched around the cue (onset of fixation cross and noise) with a 4 s pre-stimulus and a 6.5 s post stimulus interval. Artefact correction was performed in three steps: visual inspection and removal of trials with excessive artefacts occurring at all channels (on average 22.7 out of 546 trials), and marking of bad channels that were excluded from the ICA and interpolated afterwards (one each for four participants, zero for all others); removal of eye blinks and muscular artefacts by visually inspecting and then removing ICA components (rejecting on average 26.6 out of 64 components); and automatic removal of trials with activity exceeding ± 150 *µ*V (on average 25.8 trials). For the encoding models, the epoched data were low pass filtered (6th-order, two-pass butterworth filter, 25-Hz cut-off).

### Analyses of response times

All behavioral analyses were performed in R, version (version 3.3.0, R Core Team, 2016), using linear mixed effect models from the *lme4* package (Bates et al., 2015), as well as plotting functions from the *ggplot2* package (Wickham, 2009). Response times were log-transformed and the first mini-block per condition was removed from the analyses to allow participants to adapt to the new condition. Trials with response times below or above 2.5 individual standard deviations were removed as outliers. To generally assess whether hazard affects response times, we computed a linear mixed effect model with one regressor of interest (fixed effect), namely the value of the hazard function for the respective condition at the time point of the occurrence of the probe (0-centered), plus we modelled a random intercept and random slope for the effect of hazard over participants.

Second, to test the differential fit for each of the two hazard functions in each condition, we computed one linear mixed effect model per condition, using only the data from that condition and both, the monotonic and modulated hazard functions additively as fixed effect regressors, plus a random intercept and random slopes for both hazard regressors over participants.

To obtain F- and p-values for the fixed effects, we used the summary function from the *lmerTest* package (Kuznetsova et al., 2016). As an estimate of effect sizes, we report conditional and marginal *R*^2^ values (Nakagawa and Schielzeth, 2013; Johnson, 2014), which indicate the variance explained by the full model, and by the fixed effects, respectively (obtained from the *MuMIn* package, Bartoń, 2017).

### EEG Analyses: Forward Encoding Models

To test whether the time domain EEG signal tracks temporal hazard, we used a forward encoding model approach as previously described by Lalor et al. (2006; 2009), in which a time resolved regressor (like a speech envelope, or in our case the hazard function), is used to predict a time-resolved neural signal (EEG or MEG time domain or frequency data) for a range of negative and positive time lags between the two signals for each trial, using ridge (or L2-penalized) regression (Hoerl and Kennard, 1970; see also Biesmans et al., 2017; Fiedler et al., 2017; O’sullivan et al., 2015)

The approach is conceptually similar to cross-correlation, but more robust to overfitting due to usage of regularized regression. For these analyses, we used the Matlab-based multivariate temporal response function (mTRF) toolbox (Version 1.3, Crosse et al., 2016). The resulting regression weights over lags, termed *temporal response function (TRF), w(τ,n)*, can be understood as a filter with which the stimulus s(t) is convolved to obtain the response r(t,n) (Crosse et al., 2016).

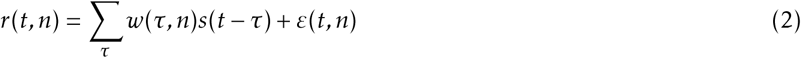

Here, 1 … T reflects sampling points of r (EEG response, measured at N channels), and s (stimulus) over time, and *τ* the sample-wise time-lag between s and r. For instance, the EEG signal might show a relation to the sensory stimulus not at lag 0 (i.e., simultaneously), but shifted in time, for example with a lag of 100 ms relative to the stimulus signal. Similar to correlation coefficients, the relation can be positive or negative. *ε*(t, n) reflects the residual errors, not explained by the model. w(*τ*,n) can be obtained using by minimizing the mean squared error, which is implemented via regularized ordinary least squares:

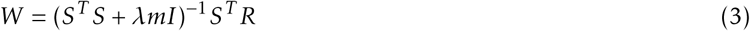

where S is the stimulus over time points t (rows), shifted over lags *τ* (columns). A regularization parameter *λ* (the ‘ridge parameter’) is introduced to prevent over-fitting. *λ* is multiplied with m, the mean of the diagonal elements of S^T^S, and with the identity matrix I, and added to the covariance matrix S^T^S (Biesmans et al., 2017). Here, we used *λ* = 4, which we had found to be the minimal value to obtain stable TRF by visually inspecting the ‘ridge trace’: the peak value of the TRF (at lag 0) over different values for *λ*. To illustrate the overall shape of the temporal response functions in an unbiased manner, before selecting specific channels and time points, we computed global field power over channels.

### EEG encoding models of auditory events

In a first step, in order to assure that the forward modelling approach works even for our relatively short epochs, we computed TRF for the encoding of the auditory events, that is, the target tones embedded in ongoing noise. To test for the encoding of the target tone onset in the EEG signal, we created for each trial a time domain stimulus vector (from 0.5 s to 3.5 s after cue-onset) by inserting the target tone into a vector of zeros, at the time point when it occurred on that trial (as in Fiedler et al., 2017). We computed stimulus envelopes by taking the absolute values of the analytic signal, then down-sampled the signal to the sampling frequency of the EEG data (200 Hz), low pass filtered (6th-order, two-pass butterworth filter, 25 Hz cut-off), and took the halfwave rectified first derivative (see Figure 3).

**Figure 3:**
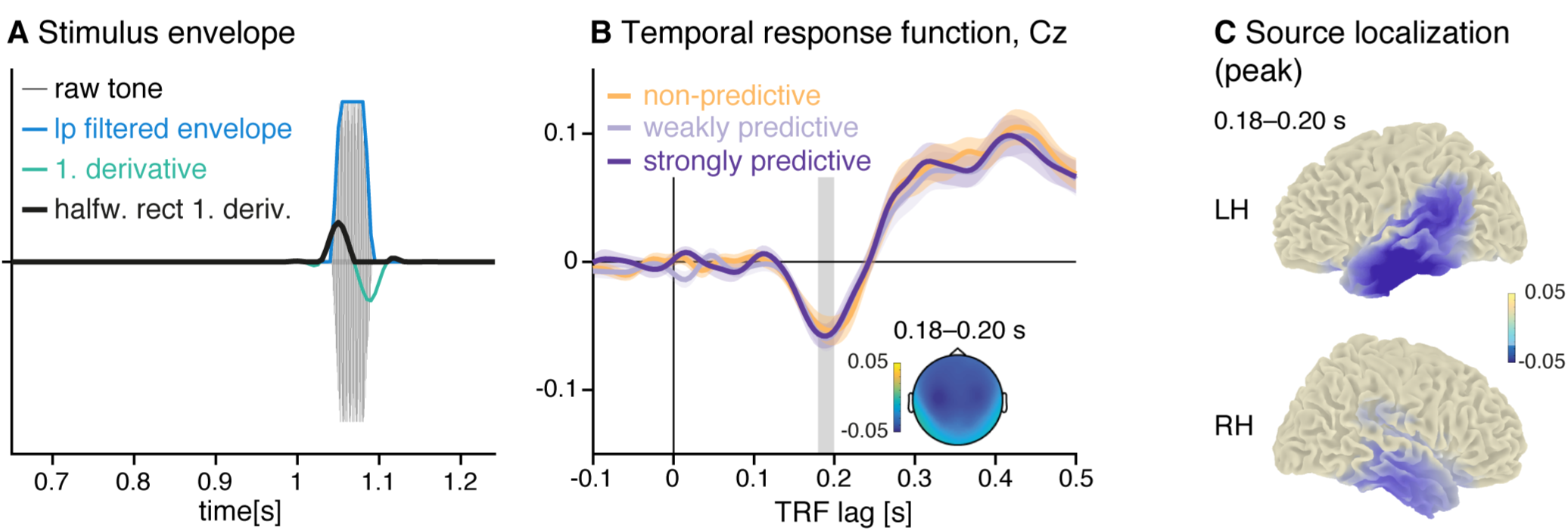
Encoding models trained on the target-onset envelopes. A: Preparation of stimulus envelopes: stimulus vectors were computed by inserting the target tone into a vector of zeros at the time of its occurrence, extracting the envelope and subsequently computing the halfwave rectified first derivative (black line). B: Temporal response functions to the target tone for all three experimental conditions. C: Topographies and source reconstructions for the negative peak of the temporal response function (averaged over conditions, 0.18–2.0 s), suggesting auditory generators.

### EEG encoding models of temporal hazard

To assess whether and how the brain tracks temporal hazard, we computed temporal response functions by training three encoding models with the temporal hazard functions as regressors (see schematic depiction in Figure 2). We trained one encoding model on the data of each of the three conditions (non-predictive, weakly predictive, and strongly predictive) by using the monotonic and modulated hazard functions jointly as regressors. This way, we obtained per encoding model two temporal response functions (TRF): one TRF reflecting the regression weights for the monotonic hazard function, and one TRF reflecting the regression weights for the modulated hazard function. Second, to test whether the EEG data from each condition encodes the temporal hazard applying to each of the three conditions, we predicted the EEG responses from each of the three models and tested the fit of the predicted with the original EEG responses from all three conditions by computing Pearson correlation coefficients between predicted and original data.

Encoding models were computed for single trials, allowing to take into account the variable intervals between cue and target. Only trials with foreperiods longer than 0.65 s were used and cut from 0.5 s after cue onset (to remove the cue-evoked activity). Importantly, we omitted the response to the target tone by setting the EEG signal to 0 after the time point of target onset on the given trial, because temporal hazard is relevant only until the occurrence of the expected event. Both, the EEG Data and hazard function vectors were low-pass filtered (using a 6th-order two-pass butterworth filter with a cut-off frequency of 25 Hz). The hazard functions (also cut from 0.5 s after cue-onset) were normalized by dividing through their maximum, and the EEG data for each trial were z-scored over the temporal dimension. The lags for the temporal response function were chosen as −0.2–0.6 s. To predict EEG data, we used a leave-one-out procedure over trials, in which we predicted EEG data for one trial, by using the average TRF from all other trials in that condition. To test whether the models are able to distinguish between the data from the three conditions, we also predicted EEG data from the models trained on the other two conditions. We thus obtained for each condition three data vectors (per trial, participant, and channel), from the models trained on the non-predictive, weakly predictive and strongly predictive conditions, respectively. Next, we computed per trial Pearson correlation coefficients between the predicted EEG data and the original data to test which of the three models provides the best fit with the original condition data (referred to as ‘testing condition’ in the figures). Correlation coefficients were Fisher-z transformed. To assess whether the correlations were different from zero, that is whether the models had any predictive power, we computed parametric 95% confidence intervals for the true correlation based on the t-distribution.

Then, to assess relative fits between the models trained on the data from the three different conditions, we computed a ‘robust’ index of fisher-z transformed correlations for each trial. To deal with different signs of individual correlation values, we applied the inverse logit transform to individual correlation values and then computed the index as the normalised difference between the correlation obtained from the model trained on the data from the non-predictive condition and the correlation obtained from training on the data from the strongly predictive condition. A correlation index larger than zero indicates a relatively better fit obtained by the model trained on the non-predictive condition, while a correlation index smaller than zero indicates a relatively better fit obtained by the model trained on the strongly predictive condition. A correlation index of zero would indicate that both models perform equally well. To test the significance of the differences in correlation indices between conditions, we performed a second-level cluster permutation test on the indices obtained for each participant over all electrodes (two-tailed, alpha 0.025, 1000 permutations).

### Source localization

To localize the peaks of the temporal response functions from the sensor-level analyses, we performed an additional source localization analysis. For the co-registration of the EEG electrode positions we used, due to the non-availability of individual anatomical models for our participants, a standard electrode layout, a standard MRI template, and a standard head model (based on the boundary-elements method) from the FieldTrip software (Oostenveld et al., 2003). Reconstruction of sources from EEG data based on template anatomical models has been successfully performed by previous studies from our own group and others (for instance Helfrich et al., 2017; Strauß et al., 2014; Praamstra et al., 2006; Bendixen et al., 2014).

Data were re-referenced to the average of all channels, and individuals’ lead field matrices were calculated with a 1-cm grid resolution. For each participant a spatial filter was computed by applying linearly constrained minimum variance (LCMV, Van Veen et al., 1997) beamformer dipole analysis to trial-averaged data (dipole orientation was not fixed). Single trials where then projected into source space and a PCA was computed on single trial three-dimensional time course data (x,y,z direction) to extract the dominant orientation (first component). Then, temporal response functions were computed on the single trial data from each source, exactly as described for the sensor level data. For visualization, the source-level peaks of the average temporal response functions over participants were interpolated and projected onto a standard MNI-brain. To illustrate the temporal response functions in supplementary motor area, we extracted single voxel activity from the labels from the AAL atlas (left and right SMA), implemented in fieldtrip (Tzourio-Mazoyer et al., 2002), and averaged over all voxels in the label.

### Control analysis on resting-state data

As a control analysis, we computed the encoding models on time domain EEG snippets from a resting-state EEG data set. We used three minutes of eyes-open resting-state recordings that had been acquired from 21 participants in an independent study (Wöstmann et al., 2015, 2014). We applied the same high- and low-pass filters to these data as described above, re-referenced to linked mastoids, down-sampled to 200 Hz, and used ICA to remove eye movements. Then, for each of the participants we randomly selected snippets from the resting-state data and replaced the original trial data by those snippets. To simulate 24 participants, we performed a second random selection for three out of the 21 resting-state participants. Encoding models were computed for these data, exactly as described for the original task data.

## Results

### Temporal hazard affects response times

To test whether response times vary with temporal hazard, we first computed an overall linear mixed-effects model jointly for all conditions with hazard as continuous regressor. As hazard, we input the value of the hazard function used for that condition at the time point of target occurrence (as depicted in Figure 1E). Response times to a given target were strongly influenced by hazard at target occurrence, shown by a significant main effect of hazard (F(1,23.1)= −4.2, p < 0.001; conditional R^2^ = 0.445, marginal R^2^ = 0.003).

Furthermore, we tested whether response times indicate a distinction between the three different hazard functions used, relating to the approach used by Janssen & Shadlen (2005) and Bueti et al. (2010). To this end, we separated the data from the three conditions and used as regressors the hazard functions from the non-predictive (monotonic hazard function) and strongly predictive conditions (modulated hazard function). Note that the weights obtained for each regressor indicate the influence of this particular regressor on the data when all other regressors are kept constant, that is, the unique variance it explains. We expected that if observers updated their temporal expectations between conditions, the relative weights obtained for the two hazard functions should differ between conditions.

For all three conditions, we obtained a negative weight for the monotonic hazard function, showing that response times decreased with rising hazard. The effect for the monotonous hazard function was significant for all conditions (marginally only in the weakly-predictive condition; p values are given in Table 1). On average the weights for the modulated hazard function were positive in the non-predictive condition, but negative in the two predictive conditions, but the modulated hazard function did not produce a significant effect in any condition (for p-values see Table 1).

**Table 1:** Linear mixed effect model of response times. Results of the three linear mixed effect models computed separately to predict response times from the non-predictive, weakly and strongly predictive conditions with the monotonic and modulated hazard functions. The first value in each cell gives the parameter estimate (i.e. the unit change in log response times by this factor when keeping all other factors constant) and its standard error in brackets. P-values indicate which hazard functions significantly correlated with the response times from the respective condition. Conditional and marginal *R*^2^ values indicate the variance explained by the full model, and by the fixed effects, respectively.

| condition | non-predictive | weakly predictive | strongly predictive |
| --- | --- | --- | --- |
| monotonic HF | **-0.14 (0.06)**<br>**$p < 0.001$** | **-0.07 (0.04)**<br>**$p = 0.06$** | **-0.11 (0.04)**<br>**$p < 0.01$** |
| modulated HF | 0.04 (0.04)<br>$p = 0.29$ | -0.03 (0.02)<br>$p = 0.18$ | -0.03 (0.02)<br>$p = 0.27$ |
| Conditional $R^2$ | 0.448 | 0.442 | 0.429 |
| Marginal $R^2$ | 0.007 | 0.001 | 0.002 |

As shown in Figure 1F, the relative contributions of the modulated and monotonic hazard functions changed over conditions, as indicated by an ANOVA computed on single subject’s ratios between the weights for the modulated and monotonic hazard functions (F(2,69) = 19.98, p < 0.0001). Post-hoc tests revealed significant differences between the non-predictive and both predictive conditions (p < 0.001, fdr-corrected).

In sum, these results show that while the monotonic hazard function affected response times in all conditions, the relative contribution of the monotonic and modulated hazard functions differed between conditions, suggesting that response times do distinguish between the different hazard functions.

### Temporal response functions of auditory stimulus encoding

To assess the neural encoding of target tones and to assure the overall applicability of forward-encoding models to these data, we trained encoding models on the time domain EEG data from all conditions, using target onset envelopes as regressors. The resulting trial-averaged temporal response functions (TRF, shown in Figure 3B) show a negative deflection around lag 180 ms, whose topography and source reconstructions suggests sources in auditory areas. The TRF showed no condition differences within the range of lags applied. This initial analysis proved that even with relatively short epochs, the approach can reveal the encoding of auditory information.

### Temporal response functions reveal the neural tracking of temporal hazard

Our main analysis sought to test whether the time domain EEG signal contains signatures of temporal hazard. We computed one encoding model per condition (per participant, trial and channel), using as regressors the monotonic and modulated hazard functions obtained from the non-predictive and strongly predictive conditions (Figure 4A). For each encoding model (i.e per experimental condition), we obtained two TRF, one for each of the two hazard functions (as shown in Figure 4B). The TRF obtained from the monotonic hazard function showed a peak at lag zero in global field power, which results from a negative deflection with a fronto-central scalp distribution in all conditions. A cluster-permutation test on the single subjects’ TRF (from −0.05 to 0.05 s) revealed a marginally significant greater negativity for the peak for the TRF trained with the data from the non-predictive compared to the predictive condition (0.035–0.05 s; p = 0.09).

**Figure 4:**
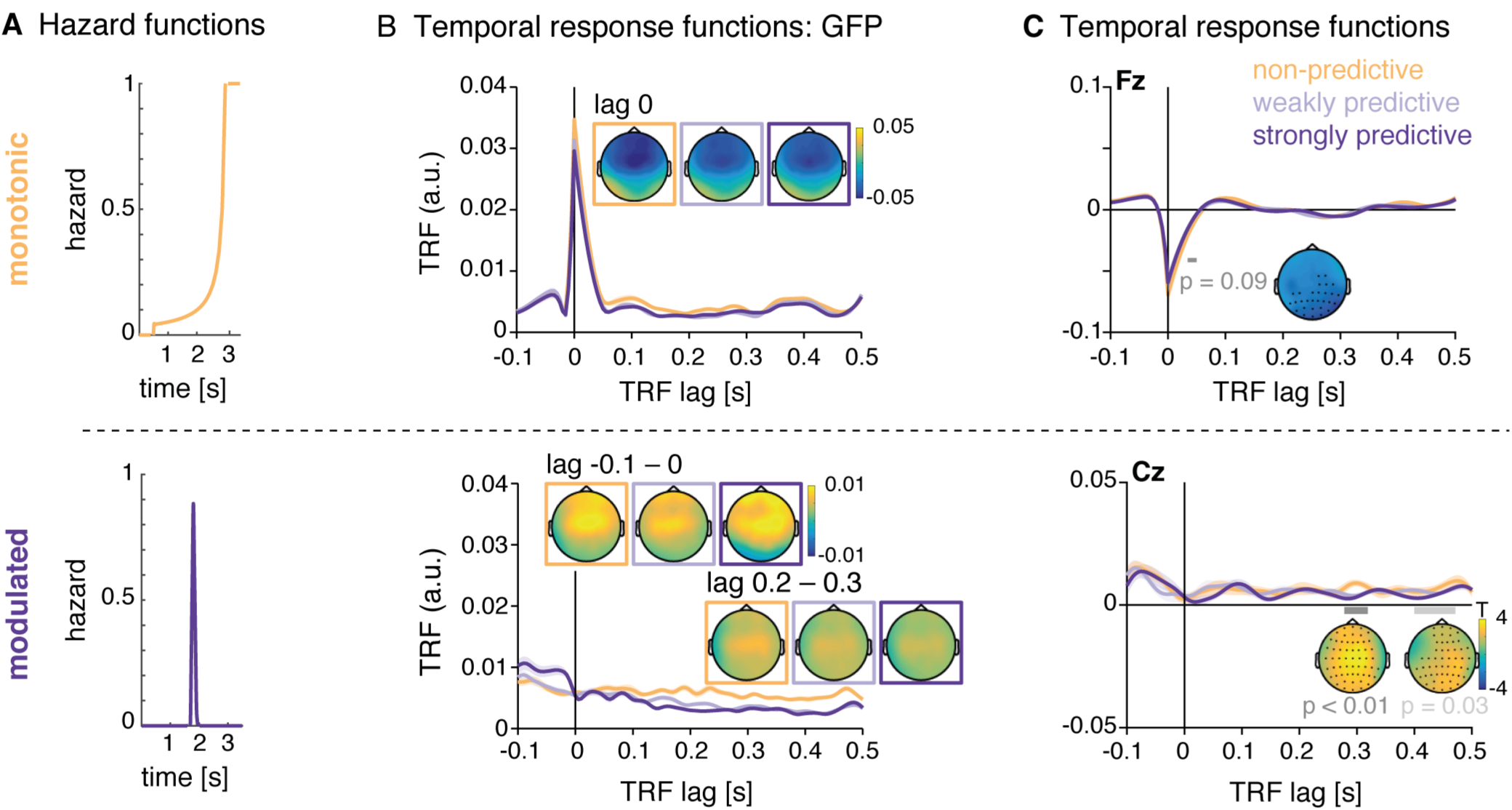
Temporal response functions (TRF) for the encoding of temporal hazard, separately for the different hazard components: A: Hazard functions, as used as regressors for the encoding models. Top: monotonic hazard function (orange); bottom: modulated hazard function (violet). B: Left: Global field power (GFP) of the TRF for the monotonic (top) and modulated (bottom) hazard function. Shaded areas indicate SEM over participants. Topographies show the scalp distribution of the TRF at lag 0 for the monotonic hazard function and lags −0.1–0 s and 0.2–0.3 s for the modulated hazard function. C: Average temporal response functions for the monotonic (top, electrode Fz) and modulated (bottom, electrode Cz) hazard function.

The TRF for the modulated hazard function showed no peak at lag zero, but, in in global field power, a stronger positivity at negative lags and a rather a sustained difference between conditions starting at a lag of about 150 ms. However, significant differences (testing for all latencies from −0.1–0.5 s) were found only for the later lags: between 0.28–0.32 s (p < 0.01) and 0.42–0.47 s (p = 0.03).

### Encoding models of temporal hazard distinguish between experimental conditions

To assess the relative fit of the three encoding models trained on the data from each condition, we predicted EEG data based on each of the three models. As a measure of prediction accuracy, we then correlated each trial of predicted EEG with the original EEG data. The predicted EEG signals averaged over trials are shown in Figure 2. As shown in Figure5A, predicted and original EEG data correlated positively, around r=0.07 on average. As shown by the confidence intervals (colored vertical bars in Figure 5A), the correlations were significantly different from zero (and from the correlations obtained with the resting-state data, grey vertical bars in Figure 5A).

**Figure 5:**
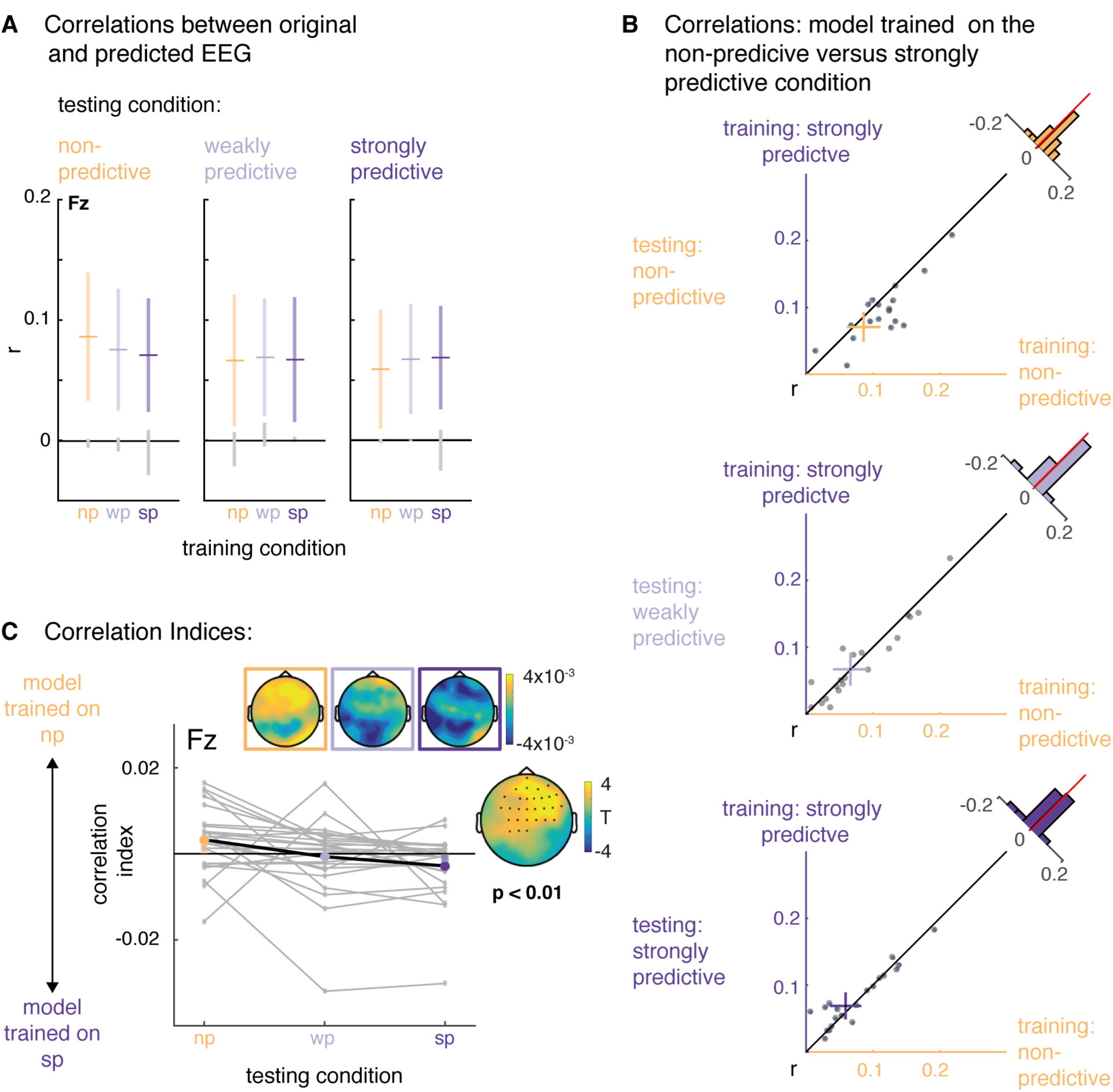
Relative model fits. A: Mean (horizontal bars) and 95% confidence intervals (colored vertical bars) for the correlation between the original EEG data from non-predictive, weakly predictive, and strongly predictive conditions (testing condition, displayed in the horizontal panels; np, wp, sp, stand for non-predictive, weakly predictive and strongly predictive) and the predicted EEG from the models trained on the non-predictive, weakly and strongly predictive conditions (on the x-axis and color-coded). The grey vertical lines display the confidence intervals obtained from the resting-state data. B: Scatterplots, displaying for each condition (‘testing condition’, vertical panels) single participants’ correlations between the data from that condition with the data predicted by the model trained on the non-predictive (x-axis) versus strongly predictive (y-axis) conditions. C: Relative correlation indices: positive and negative values indicate better fits between the original data per condition (x-axis) and the data predicted by the model trained on the non-predictive and strongly predictive condition, respectively. The insets show the topographic distribution of the indices and the T-values for the statistical comparison of the non-predictive and the strongly predictive conditions.

To assess the relative fits of the three different models trained on the data of the three conditions, we computed correlation indices contrasting for each trial the fit of the original EEG data with the predicted EEG signals from the models trained on the non-predictive versus strongly predictive conditions. The resulting index is positive when the original data fits better with the predicted data from the model trained on the non-predictive condition, and negative when the predicted data fits better with the data from the strongly predictive condition. As shown Figure 5B, for the non-predictive condition the average index showed relatively better fits for the model trained on the non-predictive condition with the original data from that condition, and respectively between the test data from and the model trained on the strongly-predictive condition. For the weakly predictive condition, the index is at around zero, indicating no conclusive evidence for either model. A two-level permutation cluster test on individual participant’s correlation indices per trial confirmed a significant difference between the correlation indices from the non-predictive versus the strongly predictive condition (p = 0.002). In sum, the differences in correlation indices revealed that the three encoding models are sensitive to condition differences in the time domain EEG data.

### Encoding of instantaneous temporal hazard in the supplemental motor area (SMA)

To assess the anatomical sources of the temporal response functions, we computed the encoding models on EEG data projected into source space (see Figure 6). This analysis revealed that the negative peaks of the TRF for the monotonic hazard functions found in the sensor space data originate in the bilateral supplementary motor areas (SMA) and the adjacent paracentral lobule (Figure 6A, left).

**Figure 6:**
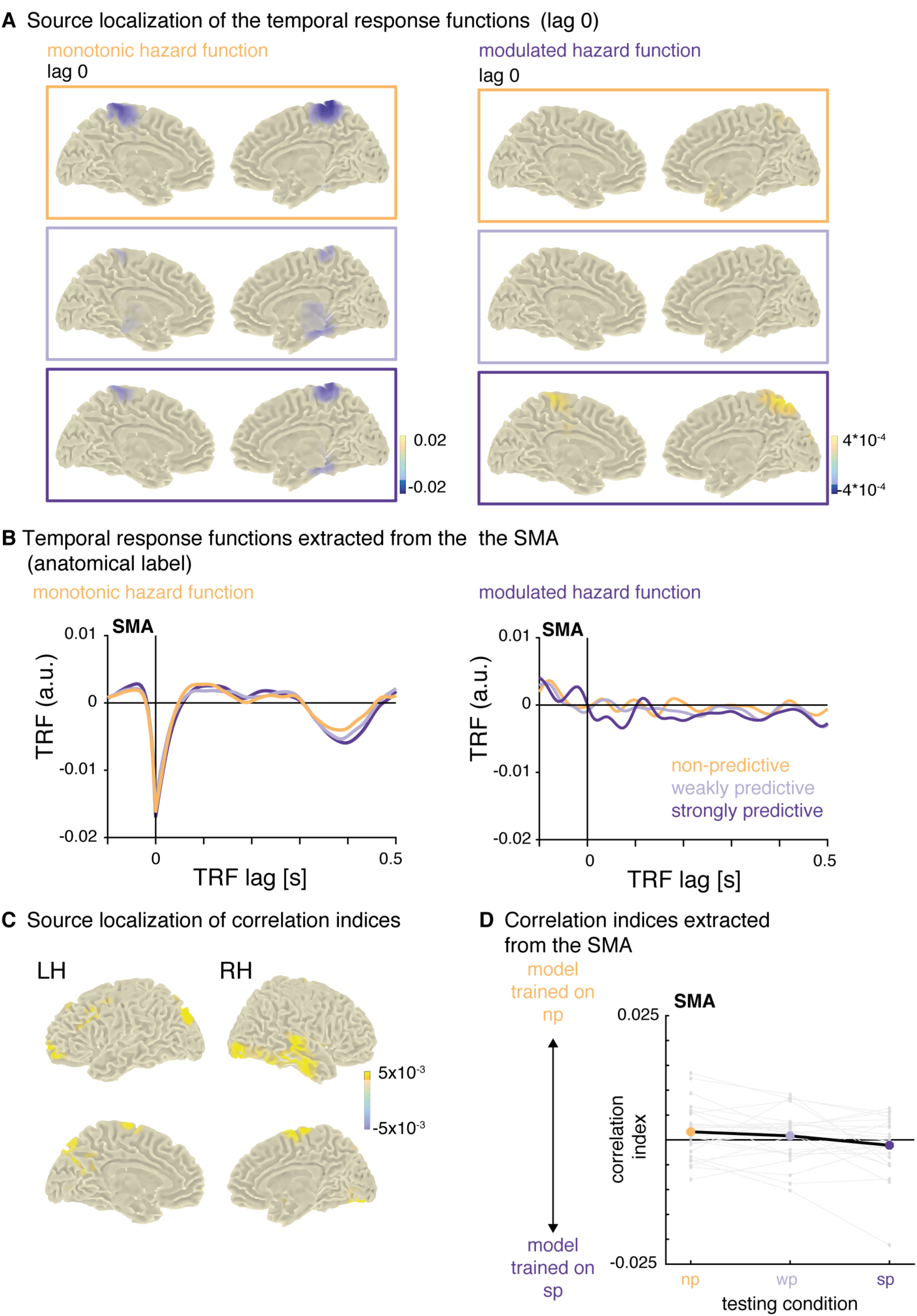
Source Reconstructions of the temporal response functions. A: Peak sources of the temporal response functions at lag 0 (the transparent overlay on the colorbar indicates the value at which the plots were masked). The strongest negative activity was localized to the bilateral supplementary motor area adjacent regions. B: Temporal response functions extracted from the bilateral supplementary motor area (SMA). C: Sources of relative correlation indices: difference between testing conditions non-predictive and strongly predictive (x-axis in panel D). D: Relative correlation indices, extracted from the bilateral SMA: positive and negative values indicate better fits between the original data per condition (x-axis) and the data predicted by the model trained on the non-predictive and strongly predictive condition, respectively.

There was no clear peak for the TRF for the modulated hazard functions in the sensor space data. In source space, the strongest activation at lag 0 was originating from the precuneus, but only in the strongly predictive condition (Figure 6A, right). To post-hoc visualize the TRF from the SMA over lags, we then extracted activity from the SMA in source-space for both hazard functions and all three conditions. The TRF for the modulated hazard function showed a stronger positivity from lags −0.05–0 s, and an adjacent negativity (lags 0–0.1 s) only for the strongly predictive condition (Figure 6C). Correlations and correlation indices computed on the data extracted from the SMA showed a similar pattern as the sensor-level data, namely better fits between the model trained on the non-predictive condition with the data from that condition, and, respectively, better fits between the model trained on the strongly predictive condition with the data from that conditions (Figure 6C,D), but the difference was not significant (T(23)=1.64, p=0.11), indicating that the SMA is not the only region contributing to the distinction between the different hazard functions. The peak sources for the positive differences in correlation indices (shown in Figure 6D) were in the bilateral SMA, precuneus and medial temporal lobes.

### Control analysis: Encoding models on resting-state data

To assess the validity and specificity of the encoding model approach, we computed the same encoding models (i.e., using the same set of hazard function regressors) on time domain EEG snippets from a resting-state data set. Figure 7 shows the temporal response functions obtained from these data. The TRF for the monotonic hazard function has a peak in global field power at lag 0 s, albeit weaker than with the original data and with much less clear topographies. The TRF for the modulated hazard function showed a sustained difference in global field power at later lags.

**Figure 7:**
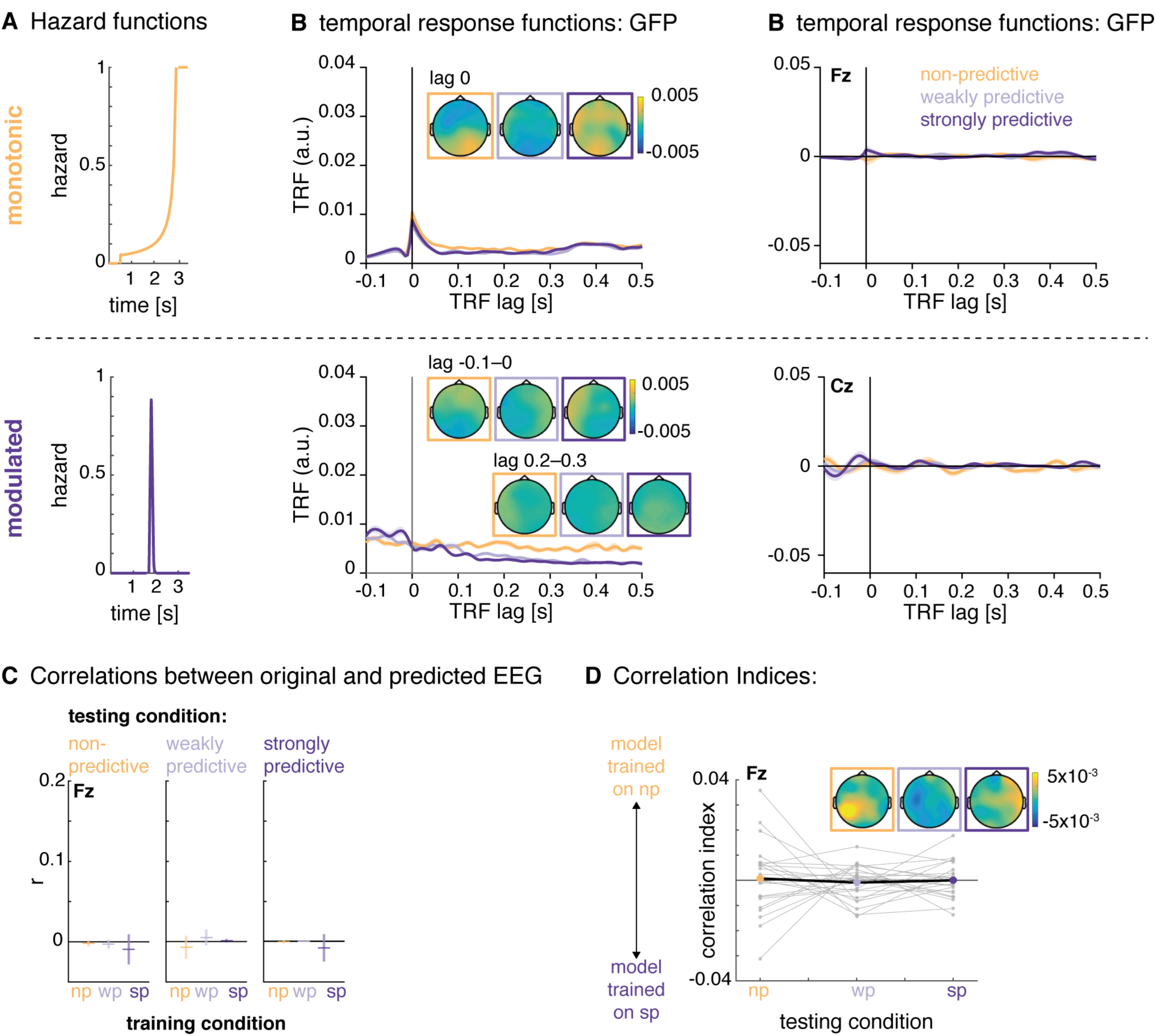
Control Analysis on resting-state data. A: Hazard functions, used as regressors for the encoding models. Top: monotonic hazard function (orange); bottom: modulated hazard function (violet). B: Global field power and average temporal response functions (electrode Fz) for the monotonic (top) and modulated (bottom) hazard function. Shaded areas indicate SEM over participants. Topographies show the scalp distribution of the TRF at lag 0 for the monotonic hazard function and lags −0.1–0 and 0.2–0.3 s for the modulated hazard function. C: Mean (horizontal bars) and confidence intervals (vertical bars) for the correlations between the original resting-state EEG data (testing condition, displayed in the horizontal panels) and the predicted EEG from the models trained on the non-predictive, weakly and strongly predictive conditions (on the x-axis and color-coded). D: Relative correlation indices: positive and negative values indicate better fits between the original data per condition (x-axis) and the data predicted by the model trained on the non-predictive and strongly predictive condition, respectively.

The – at least partial – resemblance between the TRF obtained for the original condition data and the resting-state data confirms that a forward encoding model can pick up unspecific activity, and that the regressors used for the model (here, the hazard functions) do influence the shape of the TRF to a substantial degree. Most importantly however, when predicting EEG data from the three models trained on the resting-state data and correlating the predicted and original resting-state data, the resulting correlations did not differ significantly from zero, and the correlation index did not differ between conditions (see Figure 7 C). This control analysis rules out the possibility that the results obtained with the actual task-EEG data were due to generic properties of EEG and encoding models only. In particular, one could worry that the unequal distributions of foreperiods in combination with the zeroing of the data after the end of the foreperiod would lead to differences in the model fits between conditions. However, if this was the case, it should also surface in this control analysis which it does not.

## Discussion

Here, we have shown that the human EEG tracks temporal hazard, even in a sensory task to which time is not a task-relevant dimension. Forward encoding models trained to predict the recorded EEG signal from temporal hazard were able to distinguish between experimental conditions that differed by their unfolding of temporal hazard over time. The supplementary motor area (SMA), a key region in timing and time perception, appeared as the primary source of this ‘tracking signal’ in a brain-wide search for likely generators of these encoding-model response functions. Our findings show that the mathematical concept of temporal hazard is of use to explain human behaviour in an implicit foreperiod paradigm, and that temporal hazard bears tractable signatures in the human EEG. From a methodological point of view, applying the encoding model approach to track a stimulus as abstract as temporal hazard is a novel but promising approach to study temporal processing, as it allows to map time-resolved processes to the EEG signal.

### Temporal hazard shapes response times

First evidence for the relevance of the concept of temporal hazard to cognitive processing comes from the analysis of response times of the pitch discrimination task. Response times correlated with temporal hazard: The higher the probability of a stimulus to occur was at the time point of its occurrence, the faster the response (Figure 1E, F, Table 1). This finding replicates the well-established variable-foreperiod effect (Niemi and Näätänen, 1981), which holds that longer foreperiods, genuinely associated to larger hazard, evoke shorter response times.

Furthermore, response time analyses per condition provided empirical evidence the monotonic and modulated hazard functions affected response time differentially. The relative contributions of the modulated and monotonic hazard functions to the variation of response times changed over conditions (Figure 1F): for the monotonic hazard function we obtained negative (and significant) weights for all conditions, showing that response times in all conditions decreased with higher values of the monotonic hazard function (i.e. with elapsed time). For the modulated hazard function we observed a distinction between conditions: we obtained nominally (but not significantly) positive weights for the non-predictive condition, that is an increase in response times at the peak of the modulated hazard function, which suggests that the modulated hazard function does not explain well the response times in this condition. For the two predictive conditions, we obtained negative weights that were smaller than those for the monotonic hazard function, showing that the response times decreased at the peak of the modulated hazard function, but also towards the end of the interval where the monotonic hazard function rises. Despite the dominance of the monotonic hazard function in all conditions, these results suggest that the modulated hazard function affected response times differently in the two predictive compared to the non-predictive condition. Note that, in contrast to the EEG analyses, the response time analyses used one single value from the hazard function for each trial, namely the one at target onset, not the full time-resolved hazard function as used in the EEG analysis, which might not be enough information to distinguish fully between the different levels of temporal predictability. In sum, response time in all instances is affected by rising temporal hazard due to the passage of time, and additionally captures modulated temporal hazard. This set of results lends validity to our manipulation of temporal hazard (for previous findings in this respect see Bueti et al., 2010; Janssen and Shadlen, 2005; Karlin, 1966; Todorovic et al., 2015; Tomassini et al., 2016). While previous studies had mostly varied the average foreperiod interval, the present data assert that response time also distinguishes between different degrees of temporal predictability varied around the same average foreperiod.

### Encoding models distinguish between the different conditions of temporal hazard

To test how the brain tracks temporal hazard, we applied an encoding model approach (Holdgraf et al., 2017; Lalor et al., 2006, 2009; Mesgarani et al., 2014; Naselaris et al., 2011) to track temporal hazard in human time domain EEG, recorded during an auditory foreperiod paradigm (Herbst and Obleser, 2017). The encoding models could distinguish between the different conditions of temporal hazard (see Figure 5), showing that participants utilized the implicit variations of foreperiods in the task.

Our results are broadly in line with a number of studies showing that temporal hazard shapes behaviour and neural responses in monkeys and humans, underlining the relevance of this mathematical concept in understanding cognitive processes. In monkeys, electrophysiological activity recorded from area LIP correlated with temporal hazard during the time of anticipation of the go-signal in a set-go (Janssen and Shadlen, 2005) and a temporal reproduction task (Jazayeri and Shadlen, 2015). Notably, Janssen and Shadlen found that the best fitting representation of temporal hazard and neural recordings was provided by a smoothed version of the hazard function that takes into account the scalar variability of timing (‘subjective hazard’). Applying such a subjective hazard function did not improve model fit in the present data. Probably, this is due to the relative similarity of the hazard functions we used (same mean), as well as the lower signal-to-noise ratio of human EEG compared to single-cell recordings in monkeys.

Further evidence for the relevance of temporal hazard to cognitive processing comes from studies recording fMRI from human participants, providing evidence for a representation of temporal hazard in visual sensory areas during the foreperiod interval (Bueti et al., 2010), and in the responses to the target events in the supplementary motor area (Cui et al., 2009). Due to the temporal resolution of the bold signal, these studies used much longer foreperiods. With respect to human EEG, Trillenberg et al. (2000) and Cravo et al. (2011) showed that prior to target onset, EEG activity distinguishes between different conditions of temporal hazard. Importantly, these designs used strongly discretized foreperiod distributions with very few foreperiods over all conditions, allowing to compare averaged activity over conditions, while our approach allowed us to model single trial data. All of the above-cited studies used a set-go task in which participants had to make a speeded response as soon as the go-target appeared, or even an explicit timing task (Jazayeri and Shadlen, 2015) both of which likely further promoted the use of timing strategies. In contrast, we used a sensory discrimination task, in which the temporal aspects were strictly implicit (for discussion see also Herbst and Obleser, 2017). Thus, our results show that participants extract the temporal probabilities in the different conditions and utilise temporal hazard when performing the task.

### Neural signatures of tracking temporal hazard

Importantly, and in complement to the previously published analyses of this data set, the encoding model approach allowed us to directly assess how the human brain encodes temporal hazard for each of the three conditions differing by their hazard function. From the encoding models, we obtained one temporal response function (TRF) per condition, whose deflections from zero represent a direct relationship between the EEG time domain data and the hazard functions, and can thus be interpreted as the neural correlates of temporal hazard.

The monotonic hazard function, describing rising hazard due to the passage of time, revealed the most clear cut neural signature: a near-instantaneous (i.e., zero-lag) deflection in the response function, peaking at fronto-central sensors. This negative deflection was found in the TRF trained on all three conditions, showing that hazard due the passage of time is tracked in all conditions regardless of the manipulation of temporal predictability. The sign and sources of this peak, localized to the bilateral supplementary motor area and adjacent regions, strongly argues for a relation to the well-described contingent negative variation (CNV, Walter et al., 1964) thought to emerge from the SMA. The CNV has often been described in explicit timing tasks (Herbst et al., 2014; Macar and Vidal, 2004; Macar et al., 1999; Merchant et al., 2013; Praamstra and Pope, 2007; Wiener et al., 2010). Note, however, that we did not observe a clear CNV in the ERP analysis presented in Herbst & Obleser (2017), which shows that the encoding model approach exhibits higher sensitivity.

SMA activity, or a CNV have also been described for implicit timing tasks (Akkal et al., 2004; Cui et al., 2009; Matsuzaka and Tanji, 1996). In monkey electrophysiology, ramping activity in the pre-SMA and SMA during the foreperiod was found, when the onset of a target was predictable in time (Akkal et al., 2004; Matsuzaka and Tanji, 1996). Mento et al. (2013) observed a ‘passive CNV’ in an implicit timing task, but reported premotor cortex as its primary source, rather than the SMA, which might be explained by the rhythmic oddball design used in that study (see also Mento et al., 2013). It is also important to note that neither their study nor ours used individual MRI scans for the source localization, and thus such fine grained regional divergence should be interpreted with caution. Nevertheless, our findings converge with the literature on temporal processing and provide new evidence that temporal hazard is tracked by the SMA in a strictly implicit timing task.

The sensor-space TRF for the modulated hazard function showed no clear peaks, hence did not allow to extract a distinct neural correlate of strong predictability. We found a significant difference between the TRF from the non-predictive and predictive conditions at later lags, which, however, also occurred in the TRF trained with resting state data and therefore seems to be related to the shape of the hazard function rather than the cognitive and neural processing thereof. The source space analysis of the TRF revealed activity in the precuneus around lag zero, only for the strongly predictive condition. This points towards an involvement of the precuneus in temporal prediction, in line with a previous study reporting enhanced precuneus (and default mode network) activation in a rhythmic temporal prediction task (Carvalho et al., 2016). Extracting only the data from the SMA in a post-hoc analysis revealed a positive-to-negative deflection around lag zero for the TRF of the modulated hazard function in the strongly predictive condition only (see Figure 6B, C), which tentatively suggests that the SMA also processes the modulation of temporal hazard in that condition in an instantaneous way, but might not be the dominant region to do so.

In sum, the the TRF for the monotonic hazard functions revealed a quasi-instantaneous neural correlate of tracking hazard, localized in the SMA, while the TRF for the modulated hazard functions revealed no singular feature well-defined in space and time. In sum, our findings converge with the literature on temporal processing and provide new evidence that temporal hazard is tracked by the SMA in a strictly implicit timing task. The successful distinction between conditions by the encoding models might thus result more from the decreasing applicability of the monotonic hazard function over conditions (in line with the by-trend stronger negative peak for the non-predictive condition), than from the distinct encoding of the predictive hazard function.

### Methodological relevance of the encoding model approach to abstract, time-resolved stimuli

Our results reveal the usefulness of the forward-encoding model approach applied to a seemingly abstract, mental representation such as temporal hazard. The approach has previously been applied successfully to encode the attentional processing of sensory stimuli (Ding and Simon, 2012; Fiedler et al., 2017; O’sullivan et al., 2015; Puvvada and Simon, 2017), and semantic processing of speech streams (Broderick et al., 2017; Di Liberto et al., 2018), but never to a purely ‘mental stimulus’ such as temporal hazard. Modelling EEG data with a time resolved regressor allows us to study how the brain maintains representations of complex and dynamic features such as temporal hazard. Furthermore, the approach of computing TRF for single trials can take into account only the foreperiod interval during which timing is expected to occur, between the cue and the target onset. These intervals vary over trials and conditions, which is a challenge for standard approaches of EEG data analyses involving averaging. To validate our approach, we have provided two control-level encoding models. One for the sensory stimulus from the EEG data recorded during the task to show that the encoding model approach is sensitive: TRF (Figure 3) to auditory target onsets replicated target-evoked event related potentials shown previously for these data (Herbst and Obleser, 2017). The second control-level encoding models used the temporal hazard of interest as regressor, but were trained on resting-state instead of the actual task data (Figure 7). Thus, the encoding model approach is also specific, in that it fails when applied to surrogate or unrelated data. The shapes of the TRF computed on these resting-state data show that the regressors used for the encoding model partially affect the TRF, calling for caution in the interpretation of their shape without performing a model comparison procedure.

## Conclusions

Using an implicit probabilistic foreperiod paradigm, forward-encoding models have allowed us to quantify the tracking of temporal hazard in the human EEG. Our results suggest that the neural correlates of modulated temporal hazard only partially overlap with those of monotonically rising hazard due to the passage of time. The monotonic hazard function revealed a quasi-instantaneous neural correlate of tracking hazard, while the modulated hazard function revealed no singular feature well-defined in space and time. Source localization and a set of control analyses capture the functional and anatomical specificity of these effects, with a key role for the supplementary motor area. The model-driven approach used here illustrates how implicit variations in temporal hazard bear tractable signatures in the human electroencephalogram.

### Extended Data

The Matlab and R analysis code used for the analyses of the EEG and behavioural data are available at he Open Science Framework (https://osf.io/qbhma/). The zip-archive *ExtendedData1* contains the R-scripts for the analysis of response times, Matlab-scripts for the encoding models computed on the task- and resting-state data, as well as the Matlab-scripts used for the source localization.

## Acknowledgments

This research was supported by a Max Planck Research Group grant to JO, and a DFG grant (HE 7520/1-1) to SKH. The authors are grateful to Malte Wöstmann for helpful discussions and for providing the resting-state data, and to Steven Kalinke and Maria Goedicke for help with data acquisition.

